# Survival across habitats and sizes explains ontogenetic habitat shifts in juvenile blue crabs

**DOI:** 10.1101/2025.10.07.680048

**Authors:** A. Challen Hyman, Grace S. Chiu, Michael S. Seebo, Alison Smith, Gabrielle G. Saluta, Kathleen E. Knick, Romuald N. Lipcius

**Affiliations:** WILLIAM & MARY’S BATTEN SCHOOL OF COASTAL & MARINE SCIENCES, VIRGINIA INSTITUTE OF MARINE SCIENCE, P.O. BOX 1346, GLOUCESTER POINT, VIRGINIA 23062, USA; DEPARTMENT OF STATISTICAL SCIENCES AND OPERATIONS RESEARCH, VIRGINIA COMMONWEALTH UNIVERSITY, RICHMOND, VA 23284-9026, USA; DEPARTMENT OF STATISTICS, UNIVERSITY OF WASHINGTON, SEATTLE, WA 98195-0003, USA; DEPARTMENT OF STATISTICS & ACTUARIAL SCIENCE, UNIVERSITY OF WATERLOO, ONTARIO N2L 3G1, CANADA

**Keywords:** Bayesian modeling, Nursery habitat, Seagrass, Salt marsh, Callinectes sapidus

## Abstract

Juvenile organisms often shift habitat usage during ontogeny to balance changing needs for growth and refuge, yet fine-scale habitat shifts within narrow size classes remain understudied. We conducted a manipulative tethering experiment to quantify size-specific survival of juvenile blue crabs across three habitats – seagrass meadows (seagrass), salt marsh edge (SME), and unvegetated sand flats (sand) – in the York River, Chesapeake Bay, USA. We also accounted for spatial orientation, seasonality, and turbidity. Results were analyzed using Bayesian hierarchical logistic regression models. Survival varied with crab size, habitat, and time of year, with an interaction between size and habitat. Smaller juveniles had the highest survival in seagrass, but survival increased more rapidly with size in SME and sand habitats. By 33 mm carapace width (CW), relative survival rates in seagrass and SME were statistically indistinguishable. Meanwhile, survival in sand approached that of seagrass at approximately 45 mm CW. When considered alongside previous growth rate studies, these results suggest that seagrass offers a growth–survival advantage for crabs *<* 15 – 20 mm CW, while salt marsh and sand habitats become increasingly favorable for larger juveniles due to comparable survival and faster growth. Consequently, even across a relatively small size range of the juvenile phase, size-specific patterns in survival differ substantially across structured and unstructured habitats. Our findings align with observed patterns of size-specific habitat use and underscore the complementary nursery roles of seagrass and salt marshes, reinforcing their joint importance in habitat restoration and conservation strategies for juvenile blue crabs.

## Introduction

The population dynamics of many marine and estuarine species depend on access to multiple habitats throughout their life cycle. For juveniles, the value of a particular habitat often varies with size or developmental stage due to shifting demands for growth and survival (Werner and Gilliam 1984, Dahlgren and Eggleston 2000, Lipcius et al. 2005). Juveniles typically employ life-history strategies that balance maximizing energy intake (i.e., growth) with minimizing predation risk (Werner and Gilliam 1984, Dahlgren and Eggleston 2000). However, these two goals often conflict; habitats that promote faster growth may also expose juveniles to higher predation rates (e.g., Seitz et al. 2005, Lipcius et al. 2005). As a result, habitat use tends to shift across ontogeny in response to changing resource requirements and vulnerability to predators. Early juvenile stages, being the smallest and most susceptible to predation, generally prioritize refuge over food availability (Johnston and Lipcius 2012, Johnston and Caretti 2017). With growth, juveniles can surpass the size thresholds of gape-limited predators (Nakamura et al. 2012), increasing survival and enabling access to habitats that offer greater foraging opportunities necessary for continued development.

Literature on nursery habitats has highlighted the importance of considering all critical habitats utilized by a species throughout ontogeny (Nagelkerken et al. 2015). Focusing conservation efforts on only a subset of nursery habitats risks overlooking life-stage-specific bottlenecks that can limit population sustainability. For example, conserving initial settlement habitats without protecting intermediate habitats used by larger juveniles may result in population declines despite early-stage survival. Therefore, assessing vital rates for a single size class – or aggregating across multiple size classes – provides an incomplete understanding of nursery function (Sheaves et al. 2006; 2015). To accurately identify and conserve essential nursery habitats, it is necessary to evaluate multiple juvenile size classes concurrently across candidate habitats (Nagelkerken et al. 2015, Hyman et al. 2024b).

The blue crab (*Callinectes sapidus*) is an economically important species that relies on multiple nursery habitats during its early life stages. Ingressing postlarvae (megalopae) typically settle in seagrass meadows when available, although they may also recruit to other structurally complex habitats if seagrass is limited or juvenile densities lead to high competition for space and resources (e.g., Etherington and Eggleston 2000, Etherington et al. 2003). The prevailing paradigm holds that juveniles emigrate to unstructured habitats adjacent to salt marshes between 25 and 50 mm carapace width (CW), at which point they attain a size refuge from many predators and can exploit abundant food resources in open areas (Lipcius et al. 2005). However, recent studies report elevated abundances of juveniles 16 – 25 mm CW in salt marsh habitats (Ralph 2014, Rudershausen et al. 2021, Hyman et al. 2024b) relative to seagrass, particularly near the estuarine turbidity maximum (Hyman et al. 2022; 2024b). These findings suggest that either juveniles achieve a size refuge from predation at earlier sizes than previously assumed, or that salt marshes function as important intermediate nurseries, offering increased food availability and partial refugia suitable for supporting the growth of larger juvenile crabs prior to emigration to unstructured habitat.

Ontogenetic patterns of habitat use in juvenile blue crabs may be attributed to changes in their mortality-to-growth ratios with size (e.g., Seitz et al. 2003; 2005, Lipcius et al.2005), although evidence is circumstantial, and several important questions remain. Specifically, the refuge function of salt marsh habitat for blue crabs is not well understood. Mesocosm experiments suggest salt marsh shoots provide superior refuge to juveniles relative to unstructured habitats (Johnston and Caretti 2017, Miller et al. 2023), yet field-based evidence remains sparse. For example, juvenile survival was similar between salt marshes and unstructured habitat in a fragmented marsh system in the Gulf of America (formerly Gulf of Mexico; Shakeri et al. 2020), and no comparable field studies exist to evaluate juvenile survival among multiple structured habitats in mid-Atlantic estuaries. A comprehensive understanding of ontogenetic habitat shifts requires information on juvenile abundance, growth, and survival to evaluate the relative importance of both initial and intermediate nursery habitats across the estuarine seascape (Beck et al. 2001, Nagelkerken et al. 2015).

In this study, we conducted a manipulative field experiment, as opposed to the observational study by Hyman et al. (2024b), to investigate juvenile blue crab survival across multiple nursery habitats and crab sizes. Our objective was to evaluate whether habitat- and size-specific survival rates aligned with the documented abundance patterns in juvenile blue crabs at differing sizes across seagrass, salt marsh, and unstructured sand. Our work was motivated by a limited understanding of fine-scale ontogenetic shifts in juvenile blue crabs with size, despite well-understood principles regarding ontogenetic habitat shifts in general ecology (Werner and Gilliam 1984, Dahlgren and Eggleston 2000), and the robust association between the smallest individuals and seagrass meadows (Lipcius et al. 2007). To address our objectives, we constructed multiple Bayesian hierarchical models to evaluate the effects of two structurally complex habitats – seagrass beds (herein, seagrass) and salt marsh edge (herein, SME) – as well as unstructured sand habitat (as a control; herein, sand) across the seascape of the York River, a tributary of Chesapeake Bay in the eastern United States. Our analyses focused on juveniles ranging from *≤* 10 mm to 50 mm CW.

## Methods and Materials

### Study Area

Blue crab tethering was conducted in the York River, a lower western tributary of Chesapeake Bay. This tributary features diverse habitat types and pronounced environmental gradients, such as turbidity (Hyman et al. 2024b), and supports high densities of juvenile blue crabs across a broad range of size classes (Hyman et al. 2022). These attributes make the York River an ideal natural laboratory for evaluating nursery habitat use among multiple ontogenetic stages. For the purposes of this study, the river was stratified into downriver, midriver, and upriver sections based on geomorphology and previous work (Fig. 1). The boundary between the midriver and upriver strata was based on a complementary study (Hyman et al. 2024b), which divided the river into three equally spaced segments. In contrast to Hyman et al. (2024b), we used the Coleman Bridge (Lat = 37.2421, Long = −76.5068) to delineate the downriver and midriver strata, due to significant changes in flow dynamics at this constriction point (Stockhausen and Lipcius 2003, Wood and Lipcius 2022).

**Figure 1.**
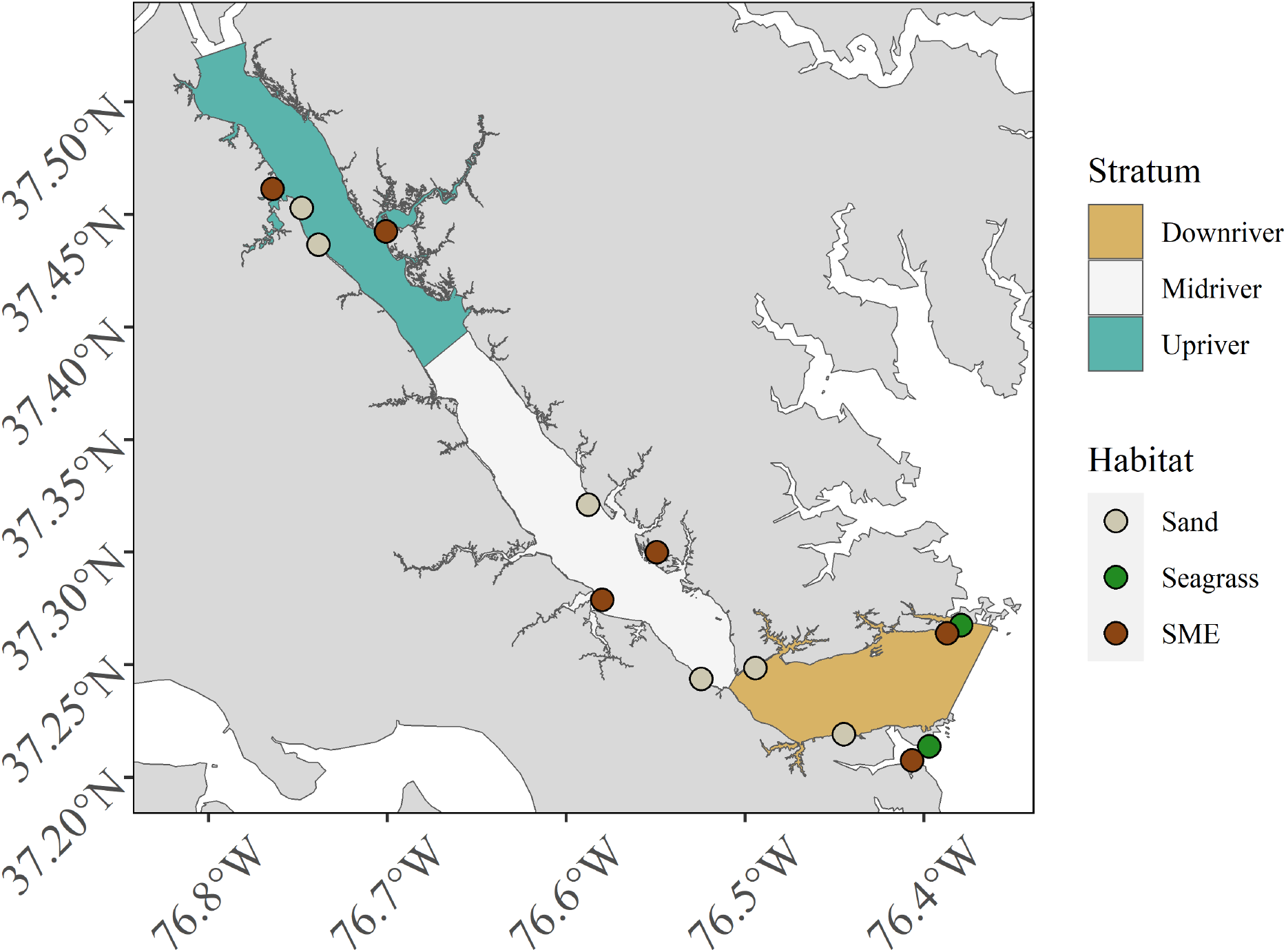
Map displaying tethering sites in the York River colored by habitat type. The York river is subdivided into downriver, midriver, and upriver spatial strata.

### Field Sampling

Sampling sites were selected using a random sampling design. Two tethering sites were randomly selected for each habitat within each stratum (n_sites_ = 14). The process included: (1) extracting geographic coordinates for the entire York River shoreline in ArcGIS, (2) filtering coordinates by habitat type and stratum, and (3) randomly choosing the predetermined number of stations for each habitat–stratum combination. Physicochemical variables salinity, temperature, and turbidity were recorded using a YSI data sonde (salinity and temperature) and a Secchi disk (proxy for the opposite of turbidity) at each site on each trip for all three sampling procedures. Within SME and seagrass, locations within the delineated habitat which were devoid of vegetation were regarded as “unstructured” and used to compare variation in survival at the patch scale. Unstructured SME habitat was defined as areas devoid of vegetation immediately adjacent to the SME, whereas unstructured seagrass habitat was defined as interstitial barren patches within or immediately adjacent to seagrass beds. In contrast, “structured” SME and seagrass habitat were defined as localities within those habitats where vegetation was present. Within SME and seagrass habitats, crabs were tethered in both structured and unstructured habitats. Only seagrass patches with 100% aerial cover were considered in structured seagrass, while sand consisted of only unstructured habitat.

Tethering was conducted at biweekly intervals between April and November, 2021 with juvenile crabs of 6 – 50 mm CW (n = 848), using an established tethering technique to assess survival (see Lipcius et al. 2005, for details). July was not considered because logistical issues prevented sampling in that month. Tethering involved attaching 30 cm monofilament fishing line to the crab’s carapace with cyanoacrylate glue. The other end of the line was tied to a stake pushed into the sediment. The stake was tied to a location pole 1 m away to minimize the effects of artificial structures that could attract predators to the tethered crab. Tethered crabs were allowed to acclimate in laboratory aquaria for 24 h prior to placement in the field.

Tethering was conducted in *<* 1 m mean low water to limit the influence of depth (e.g. Ruiz et al. 1993). At each site, five crabs were haphazardly selected and tethered in both structured (where present) and unstructured habitats. Within a habitat-structure combination, individual tethers were haphazardly spaced *∼*5 m apart. The size (CW) of each crab was measured to the nearest 0.1 mm using calipers prior to deployment, and deployed for *∼*24 h.

Molting was evident if only the exoskeleton remained on the monofilament line and was used to distinguish predation from molting in field experiments. Molts were subsequently recorded and excluded from analysis (*<* 10% of instances).

Prior to field experiments, pilot experiments were used to determine the probability of escape (i.e., crabs un-tethering themselves) and potential changes in behavior. In April, 2021 (i.e., when predation is negligible; e.g., Ruiz et al. 1993, Moody 1994), five crabs were tethered in two locations (n = 10) and checked daily. In the absence of predation, juvenile crabs remained on tethers for over a week and were able to bury themselves in sediment. We replicated this procedure in lab conditions with 10 additional crabs observed daily for 10 d. Here, only one crab escaped its tether on day eight. Since crabs in our field experiments were deployed for approximately 24 h, we concluded that the number of crabs removed due to escape was negligible.

Tethering can introduce treatment-specific bias in survival (Peterson and Black 1994, Baker and Waltham 2020). For example, tethered crabs may experience lower survival in structurally complex habitats such as seagrass and SME as a result of entanglement, but may not experience the same reduction in survival in sand, such that relative survival rates would not be comparable between these habitats. Alternatively, variation in escape behaviors (e.g., crypsis in structurally complex habitat vs. fleeing in sand) may also introduce bias. Extensive work from previous studies examining treatment-specific biases of tethering juvenile crabs in seagrass, SME, and sand has not found interactions between tethering and habitat (Pile et al. 1996, Hovel and Lipcius 2001, Lipcius et al. 2005, Miller et al. 2023); therefore, we assumed there was no treatment-specific bias in our experiments, which used similar tethering methods and habitats as those in previous studies.

### Predictors of relative survival

We evaluated multiple environmental and biological variables (hereafter, “predictors”) as potential determinants of relative survival in juvenile blue crabs. Below, we justify the inclusion of each predictor in our survival models.

Our primary objective was to assess nursery habitat quality based on relative juvenile blue crab survival. Structurally complex habitats are known to enhance refuge availability for small prey by offering a high density of interstitial spaces among biogenic structures such as seagrass shoots and rhizomes (Hovel and Lipcius 2001, Orth and van Montfrans 2002, Johnston and Lipcius 2012, Miller et al. 2023). Accordingly, we hypothesized higher survival in structurally complex habitats – specifically, seagrass meadows and salt marsh edge (SME) – compared to unvegetated sand flats (Long et al. 2013, Ajemian et al. 2015, Bromilow and Lipcius 2017). The effectiveness of structural refuge may vary across habitats. For example, seagrass shoots often provide smaller interstitial spaces that may better protect smaller juveniles than the larger spaces found among salt marsh vegetation (Orth and van Montfrans 2002). Habitat structure also varies spatially due to patchiness and other local features (Hovel and Lipcius 2001; 2002, Lipcius et al. 2005). To account for this variability, we considered two habitat classification schemes. The first, denoted “habitat”, treated all unstructured habitats as equivalent (“sand”), regardless of their spatial context. The second, denoted “habitat-structure”, distinguished between unstructured areas adjacent to seagrass (“unstructured seagrass”) or SME (“unstructured SME”) and those isolated from structured habitats, while also separately modeling structured seagrass and SME habitats.

As juvenile blue crabs grow, they are less susceptible to predation as their carapace widens and hardens, spines become more prominent, and aggressive behavior intensifies (Hines and Ruiz 1995, Hovel and Lipcius 2002, Lipcius et al. 2007, Bromilow and Lipcius 2017). Smaller juveniles are more vulnerable to predation, whereas larger individuals may attain a size refuge from some predators (Lipcius et al. 2005). Therefore, we included size as a continuous covariate in all models. Previous studies also suggest that survival may depend on interactions between habitat and crab size (Pile et al. 1996, Johnston and Lipcius 2012), so we included habitat-by-size interaction terms in a subset of candidate models.

Turbidity may reduce predation risk by impairing visual predator efficiency and decreasing light intensity (O’Brien et al. 1976, Cyrus and Blaber 1987, Howson et al. 2022, Horodysky et al. 2010, Marley et al. 2020). Along the estuarine gradient, upriver unstructured habitats tend to be more turbid than those downriver and consequently may function as effective nurseries despite lacking structural complexity (Lipcius et al. 2005, Seitz et al. 2005). We included turbidity as (*−* 1) *×* (Secchi-disk depth) as a continuous predictor of blue crab survival.

Predator assemblages can vary spatially along the river axis, potentially confounding habitat effects due to unmeasured differences in predation pressure (Posey et al. 2005, Lipcius et al. 2005). To address this, we included stratum (i.e., location along the river: downriver, midriver, or upriver) as a categorical fixed effect in all models.

Finally, predation pressure on juvenile blue crabs is known to vary seasonally, with survival typically peaking in spring and late fall and declining during summer months (Hines and Ruiz 1995, Moody 1994, Hovel and Lipcius 2001, Facendola 2010). To account for these nonlinear seasonal trends in survival, we incorporated harmonic regression terms—specifically, continuous sine and cosine functions with an annual periodicity (365 days). This approach is commonly used in time-series analysis to model seasonal effects (Hyndman and Athanasopoulos 2018, Trudeau et al. 2022, Hyman et al. 2024a; 2025). For each tether deployment, we applied sinusoidal and cosinusoidal transformations to the corresponding Julian day. For example, the sine term was calculated as: 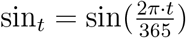 where *t* is the date of deployment, converted to Julian day. These harmonic terms allowed us to flexibly model cyclical seasonal variation in survival across the study period.

### Analysis

All data analyses, transformations, and visualizations were completed using the R programming language for statistical computing (R Core Team 2022) and the Stan probabilistic programming language for Bayesian statistical modeling (Stan Development Team 2020; 2022). Crab survival, recorded as 1 (alive) or 0 (eaten), was analyzed for the probability of survival using a hierarchical logistic regression mixed-effects model. For the *j*^th^ tether trial in the *s*^th^ site on date *t*, the model for juvenile blue crab survival is expressed as:

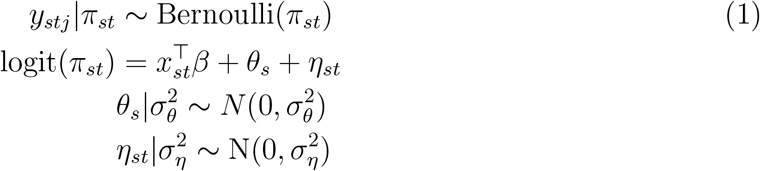

Specifically, the response, binary juvenile crab survival *y*_*stj*_ for the *j*^th^ tethering trial, is distributed as a Bernoulli random variable with a probability of survival, *π*_*st*_. Meanwhile, 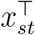 corresponds to environmental predictors at site *s* on date *t* (see Table 1 below for specific predictor combinations). Tethering trials entailed repeatedly using the same sites, which may introduce site-specific bias. Moreover, 5 m spacing of individual tethers within a site may not have been sufficient to guarantee independence (Baker and Waltham 2020). Hence, *θ*_*s*_ and *η*_*st*_ denote site-specific and site within date-specific random effects, respectively.

**Table 1.**
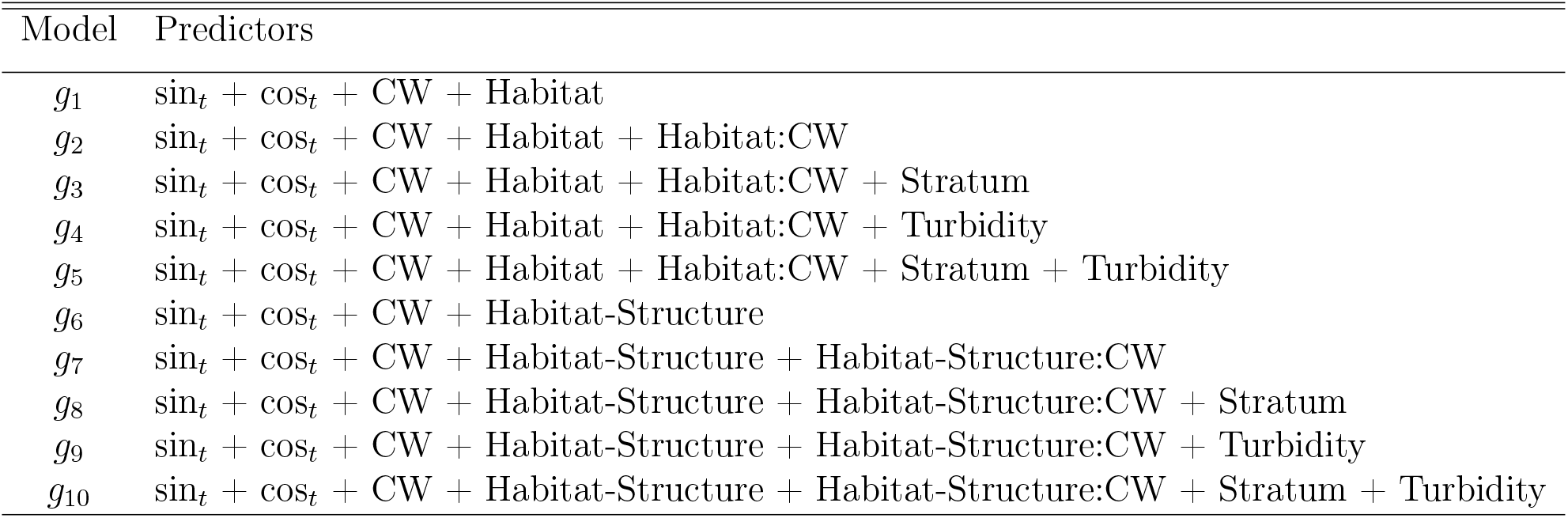
Fixed-effects predictors for the ten candidate models. **sin**_***t***_ and **cos**_***t***_ denote harmonic terms capturing seasonal effects. **CW** is carapace width (crab size); **Habitat** denotes simple habitat classifications (sand, seagrass and SME); **Habitat-structure** denotes complex habitat classifications (unstructured sand, unstructured seagrass, unstructured SME, structured seagrass, structured SME); **Stratum** denotes river strata (downriver, midriver, or and upriver). **Turbidity** is water cloudiness. A “+” denotes additive, independent effects, while a colon (e.g., Habitat:CW) denotes an interaction, indicating size effects within specific habitats.

| Model | Predictors |
| --- | --- |
| $g_1$ | $\sin_t + \cos_t + \text{CW} + \text{Habitat}$ |
| $g_2$ | $\sin_t + \cos_t + \text{CW} + \text{Habitat} + \text{Habitat:CW}$ |
| $g_3$ | $\sin_t + \cos_t + \text{CW} + \text{Habitat} + \text{Habitat:CW} + \text{Stratum}$ |
| $g_4$ | $\sin_t + \cos_t + \text{CW} + \text{Habitat} + \text{Habitat:CW} + \text{Turbidity}$ |
| $g_5$ | $\sin_t + \cos_t + \text{CW} + \text{Habitat} + \text{Habitat:CW} + \text{Stratum} + \text{Turbidity}$ |
| $g_6$ | $\sin_t + \cos_t + \text{CW} + \text{Habitat-Structure}$ |
| $g_7$ | $\sin_t + \cos_t + \text{CW} + \text{Habitat-Structure} + \text{Habitat-Structure:CW}$ |
| $g_8$ | $\sin_t + \cos_t + \text{CW} + \text{Habitat-Structure} + \text{Habitat-Structure:CW} + \text{Stratum}$ |
| $g_9$ | $\sin_t + \cos_t + \text{CW} + \text{Habitat-Structure} + \text{Habitat-Structure:CW} + \text{Turbidity}$ |
| $g_{10}$ | $\sin_t + \cos_t + \text{CW} + \text{Habitat-Structure} + \text{Habitat-Structure:CW} + \text{Stratum} + \text{Turbidity}$ |

We specified informative prior distributions for predictor coefficients (*β*) based on expert knowledge and results from prior tethering studies conducted in the same or nearby study locations. The baseline effect of unstructured sand habitat was assigned a weakly informative *N* (0, 0.5) prior, where the given parameter values denote the mean and standard deviation, respectively. Although previous studies suggest that juveniles tethered in sand exhibit the lowest survival among habitat types (Pile et al. 1996, Hovel and Fonseca 2005, Johnston and Lipcius 2012, Bromilow and Lipcius 2017), we avoided imposing a strongly negative prior due to variation in reported effect sizes (e.g., Johnston and Lipcius 2012, Bromilow and Lipcius 2017) and confounding factors such as seasonal timing—many prior studies were conducted during midsummer (July–August), when predation pressure is typically highest (Moody 2003).

For the effect of structurally complex seagrass relative to sand, we assigned a prior of *N* (2, 0.5) to reflect the strong refuge function of submerged aquatic vegetation (SAV) documented in the literature (Pile et al. 1996, Hovel and Fonseca 2005, Johnston and Lipcius 2012, Bromilow and Lipcius 2017). In contrast, the effect of structurally complex SME relative to sand was assigned a weakly informative *N* (0, 1) prior, acknowledging mixed results from mesocosm experiments that found survival in marsh habitats varied depending on predator identity (e.g., Orth and van Montfrans 2002, Johnston and Caretti 2017, Miller et al. 2023).

We also assigned an informative prior of *N* (0.05, 0.05) for the effect of CW in unstructured sand, reflecting the range of size-based survival trends reported in earlier work (Hovel and Fonseca 2005, Johnston and Lipcius 2012, Bromilow and Lipcius 2017). For interaction effects between size and habitat (relative to the size effect in sand), we used *N* ( *−*0.05, 0.05) for seagrass and *N* (0, 1) for SME. The prior for the seagrass effect reflects findings that survival is constant or may decline with increasing size in SAV (Pile et al. 1996, Johnston and Lipcius 2012), while a default weakly informative prior was used for the SME effect due to a lack of robust, size-specific survival estimates in marsh habitats. All remaining predictor coefficients were assigned default *N* (0, 1) priors, representing weakly informative, centered distributions on the logit scale. Finally, variances for random effects (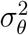 and 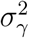) were assigned weakly informative inverse-gamma(1, 1) priors to ensure positivity and allow a broad range of plausible values while providing mild regularization.

### Model implementation and validation

We implemented the above models using the brms package (Bürkner 2017), which compiles and fits regression models using the Stan programming language for Bayesian inference (Gelman et al. 2015). For each model, we ran four parallel Markov chains, each with 5,000 iterations for the warm-up/adaptive phase, and another 5,000 iterations as posterior samples (i.e., 20,000 draws in total for posterior inference). We assessed chain convergence via inspection of trace plots and the split 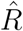 statistic to ensure values were less than 1.01, suggesting chain convergence (Gelman et al. 2015). Covariates and interactions whose regression coefficients had Bayesian confidence intervals (CIs) that excluded 0 at a confidence level of 80% were considered scientifically relevant to juvenile blue crab survival following the logic established in Hyman et al. (2024b). All CIs referenced here are highest posterior density intervals.

#### Model selection

We constructed ten candidate models to predict variation in juvenile blue crab survival. Our first five models included simple habitat characterizations—structurally complex seagrass, structurally complex SME, and unstructured sand habitat regardless of location—whereas the second five models considered habitat–structure interactions (Table 1). For our simplest models (*g*_1_ and *g*_6_), we posited that survival would be explained by an additive function of seasonality, habitat, and size. Next, we considered the effects of seasonality, habitat, size, and a habitat–size interaction on survival (*g*_2_ and *g*_7_). For the third model structure, we included seasonality, habitat, size, habitat–size interaction, and spatial stratum on survival (*g*_3_ and *g*_8_). Our fourth model structure (*g*_4_ and *g*_9_) included seasonality, habitat, size, habitat–size interaction, and turbidity, while our most complex model structure included seasonality, habitat, size, habitat–size interaction, spatial stratum, and turbidity (*g*_5_ and *g*_10_). These model structures were designed to span a gradient from simple to more complex hypotheses about the ecological drivers of juvenile blue crab survival, while avoiding over-parameterization deemed unnecessary a priori. By organizing the model set in parallel groups—one emphasizing broad habitat categories and another including habitat–structure interactions—we could compare support for alternative habitat classification schemes while systematically evaluating the additional contributions of spatial and environmental covariates.

We employed estimated log-pointwise predictive density (ELPD) and corresponding Δ_*ELPD*_ values to assess the predictive performance of each model within our set of statistical models *g*_*i*_ (Vehtari et al. 2017). ELPD values are commonly used to evaluate out-of-sample predictive accuracy, while Δ_*ELPD*_ values represent the difference between the ELPD of a given model and that of the best-performing model in the set. The ELPD and Δ_*ELPD*_ values were estimated using the approximate Leave-One-Out Information Criterion (LOO-IC; Gelman et al. 2015, Vehtari et al. 2017), with the calculations performed through the loo package in R (Vehtari et al. 2022). When two models showed similar Δ_*ELPD*_ values (i.e., *≤*4; Sivula et al. 2020), the simpler model was selected for parsimony.

### Conditional effects

To conceptualize how predictors influenced juvenile blue crab relative survival, we developed a counterfactual matrix of simulated scenarios, denoted *X*^*∗*^, where each row *i*(ie., 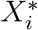) represents a unique combination of plausible values for relevant predictors. Using the coefficient posterior distributions from our best-performing model (presented in the Results section), for each set of predictors 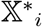 and posterior draw *d*, we computed the corresponding approximate conditional probability of survival 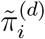 as

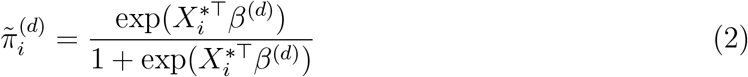

where the notation “∼ “above *π* denotes the condition of fixing the random effects for site (*θ*_*s*_) and date within site (*η*_*st*_) at 0. For our counterfactual design matrix, we considered all possible combinations of Julian day (later transformed into sin_*t*_ and cos_*t*_ terms) fixed at the first of April, May, June, August, September, and October, CW (5 to 50 mm, by 1 mm increments), and all categorical terms from our best-performing model (i.e., 828 combinations in total).

To investigate how changes in habitat may occur with size, we developed hypothetical mortality-growth ratios as a function of habitat and CW using conditional expected daily relative survival probabilities 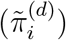 from our model and habitat-specific growth rates (denoted 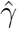) using estimates from Seitz et al. (2005). Specifically, we used the published mean aggregated blue crab growth for seagrass and upriver unstructured habitat adjacent to salt marshes from caging studies carried out between 2000 and 2001, presented on the log_10_-scale, and converted them to the natural scale (i.e., absolute CW increase). We subsequently divided these estimates by 90 days (the duration in the study over which growth was hypothesized to occur due to water temperatures and blue crab physiology) to obtain an approximate growth rate,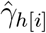, for each habitat *h* in row *i*. To convert daily conditional relative probability of survival to a mortality rate *µ*, we employed the following transformation: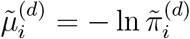 (Ricker 1975, Beverton and Holt 2012). Finally, to obtain approximate mortality-to-growth ratios, we divided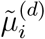 by the approximate growth rate 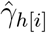. As we did not conduct the habitat-specific growth experiments, and only used mean growth rates in our approximate mortality-to-growth ratios, we stress that the habitat-specific trends in these ratios with changing size should be taken as *ad hoc* and do not reflect the full scale of uncertainty associated with growth and survival rates.

## Results

The best-performing model, as identified by LOO-IC, was *g*_9_, which described relative juvenile blue crab survival as a function of seasonal harmonic terms, habitat-structure interacting with CW, and turbidity. However, models *g*_10_, *g*_7_, *g*_8_, *g*_2_, and *g*_4_ demonstrated similar predictive performance, with Δ_*ELPD*_ *≤* 4 (Table 2); model *g*_2_ being the simplest among the six. Consequently, we selected model *g*_2_, which considered seasonality, habitat, CW, and habitat interacting with CW, for statistical inference under the principle of parsimony. Herein, all inferences are based on model *g*_2_.

**Table 2.** Model selection results from ten Bayesian logistic regression models (*g*_*i*_) predicting juvenile blue crab survival. Models are presented in order of predictive power based on the **LOO-IC**: the approximate Leave-One-Out Information Criterion. Associated metrics are **ELPD**_**LOO**_: the estimated log-pointwise density calculated from LOO; **Δ**_**ELPD**_: the relative difference between the ELPD of any model *j* and the best model in the set; **p**_***loo***_: the effective number of parameters. The selected model (*g*_2_) values are presented in bold font.

| Model | LOO-IC | ELPD<sub>LOO</sub> | $\Delta_{\text{ELPD}_{\text{LOO}}}$ | $p_{\text{LOO}}$ |
| --- | --- | --- | --- | --- |
| $g_9$ | 960.60 | -480.30 | 0.00 | 77.22 |
| $g_{10}$ | 961.17 | -480.58 | -0.28 | 76.18 |
| $g_7$ | 961.39 | -480.69 | -0.40 | 76.94 |
| $g_8$ | 963.60 | -481.80 | -1.50 | 78.17 |
| **$g_2$** | **965.90** | **-482.95** | **-2.65** | **73.81** |
| $g_4$ | 967.68 | -483.84 | -3.54 | 77.13 |
| $g_5$ | 969.38 | -484.69 | -4.39 | 76.77 |
| $g_6$ | 971.80 | -485.90 | -5.60 | 73.81 |
| $g_3$ | 972.02 | -486.01 | -5.71 | 78.53 |
| $g_1$ | 972.09 | -486.05 | -5.75 | 73.82 |

The posterior distributions of all predictor coefficients in model *g*_2_ excluded 0 from their 80% CIs, indicating all predictors meaningfully explained variation in juvenile blue crab relative survival (Table 3). Both seagrass and SME conferred higher survival to small juveniles relative to sand. Survival in sand habitat (the reference level) increased with crab size, although survival increased more rapidly in SME and more gradually in seagrass relative to sand. Both sin_*t*_ and cos_*t*_ also exhibited strong influences on survival, indicating that seasonality played an important role in relative survival over time.

**Table 3.** Posterior summary statistics – maximum *a posteriori* (MAP) and 80% CI – the model intercept and predictor coefficients from our best-performing model (*g*_2_). The intercept refers to the reference habitat level (sand). Terms **sin**_***t***_ and **cos**_***t***_ denote harmonic predictors capturing seasonal effects. Terms **SG** and **SME** are the effects of seagrass and salt marsh edge habitats relative to sand, respectively. **CW** is carapace width (crab size) in sand habitat, while **SG:CW** and **SME:CW** are the effects of carapace within seagrass and SME relative to sand, respectively. All values are rounded to two decimal places.

| Predictor | 10% | MAP | 90% |
| --- | --- | --- | --- |
| Intercept | -0.95 | -0.43 | 0.09 |
| $\sin_t$ | 0.86 | 1.17 | 1.52 |
| $\cos_t$ | 1.04 | 1.64 | 2.20 |
| SG | 2.03 | 2.50 | 2.97 |
| SME | 0.39 | 0.79 | 1.27 |
| CW | 0.03 | 0.04 | 0.06 |
| SG:CW | -0.05 | -0.03 | -0.01 |
| SME:CW | >0.00* | 0.02 | 0.04 |

\* The precise value is 0.0046.

Conditional effects plots revealed complex, size-dependent interactions between habitat type and survival across seasons. Survival followed a nonlinear temporal trend, peaking in spring (highest in April), declining through summer (lowest in August), and rising again in fall (Fig. 2). For the smallest crabs, relative survival was highest in seagrass, intermediate in SME, and lowest in sand. However, the influence of size on survival varied by habitat: survival increased more steeply with size in SME and sand than in seagrass. By 33 mm CW, the *maximum a posteriori* (MAP) values of 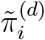 (i.e., approximate conditional probability of survival) in seagrass and SME converged. At 50 mm CW, the 80% credible intervals for all three habitats overlapped, indicating that survival differences among habitats diminished with increasing crab size.

**Figure 2.**
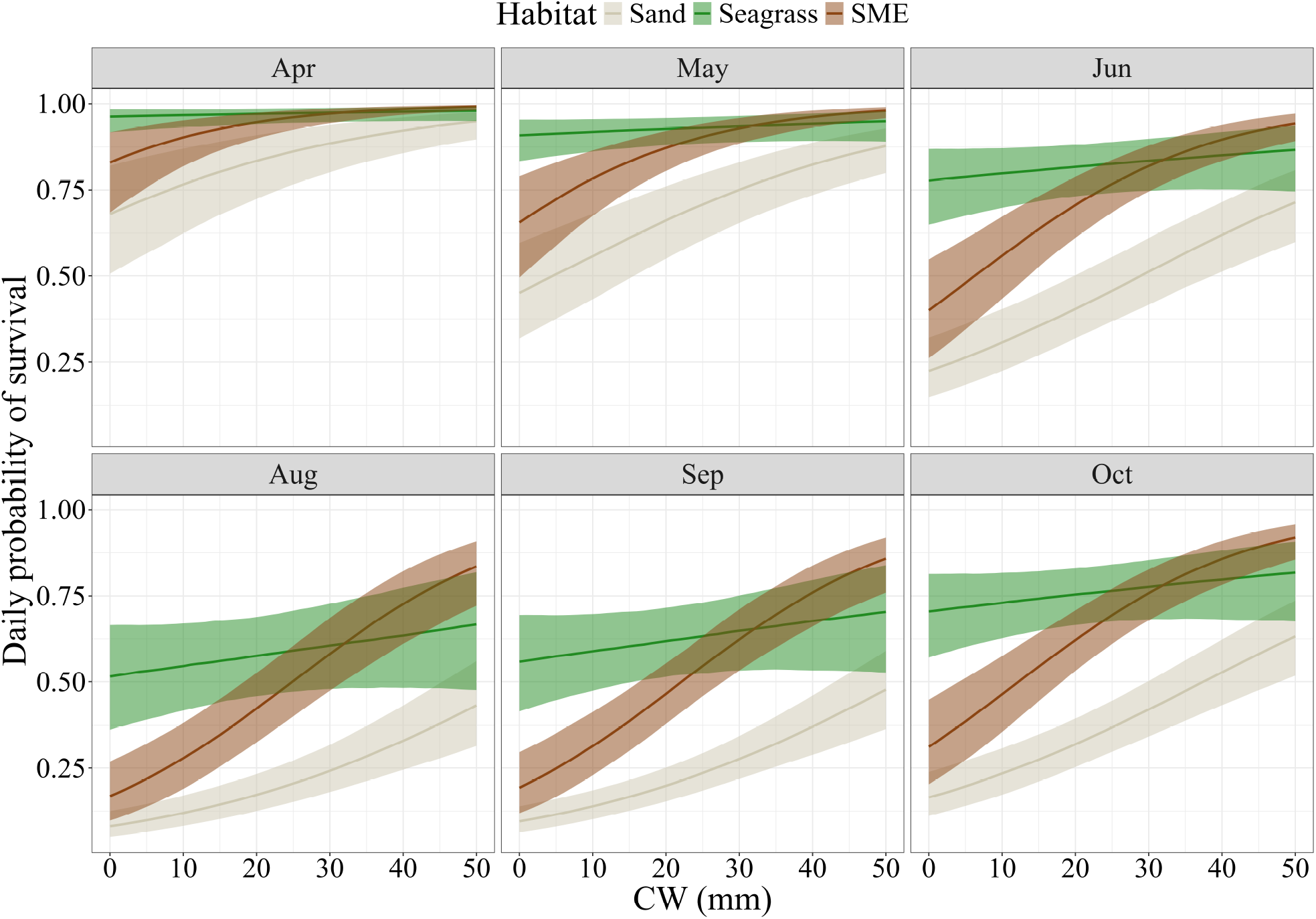
Approximate conditional posterior probability of survival 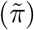 of juvenile blue crabs as a function of habitat, carapace width (CW), and time of year. Colored lines represent maximum *a posteriori* values; shaded bands indicate 80% pointwise credible intervals. Each panel corresponds to the first day of a different month within the study period, illustrating seasonal variation in predicted conditional survival across habitats and body sizes. See Hyman et al. (2022) for details on the construction of such plots.

At smaller crab sizes, approximate conditional relative mortality-to-growth ratios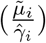 were lowest in seagrass, intermediate in salt marsh edge (SME), and highest in sand (Fig.3). However, uncertainty in these quantities was substantial, with overlapping credible intervals between seagrass and SME. At 22 mm CW, the MAP values of conditional mortality-to-growth ratios in seagrass and SME intersected, after which ratios declined more rapidly in SME. As crabs approached 50 mm CW, the conditional mortality-to-growth ratio was generally lower in SME than in seagrass. By 33 mm CW, the 80% CIs for ratios in seagrass and sand overlapped, suggesting that trade-offs between growth and conditional survival became comparable in these two habitats.

**Figure 3.**
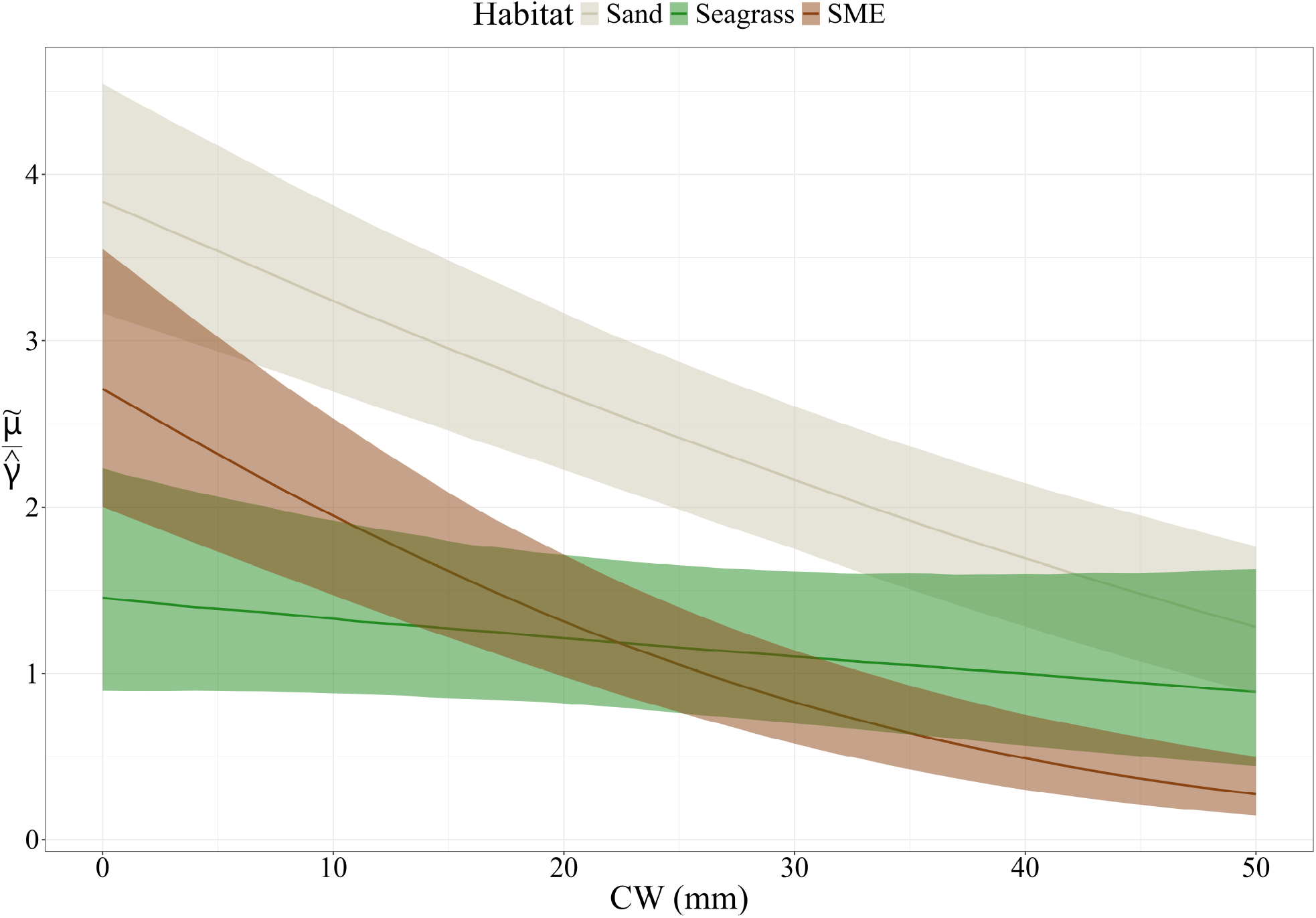
Conditional relative ratios of mortality 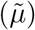 to growth (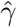, regarded as fixed) of juvenile blue crabs as a function of habitat and carapace width (CW) in August. Colored lines represent maximum *a posteriori* values; shaded bands indicate 80% credible intervals.

## Discussion

For a given life stage, habitat-specific vital rates are key determinants of a habitat’s relative importance within the broader seascape. Accurately estimating size- and habitat-specific production requires understanding the biotic and abiotic factors that drive these rates both within and among habitats. In marine nursery systems, studies of survival, growth, abundance, and ontogenetic habitat shifts have largely focused on broad size classes across the juvenile phase. However, fine-scale ontogenetic shifts in habitat use can occur over much narrower size ranges – patterns that remain understudied but are critical for the effective conservation and restoration of nursery habitats.

In our study, structurally complex seagrass meadows provided the highest survival for newly settled juvenile blue crabs, followed by structurally complex salt marsh edge (SME) habitat and unstructured sand. These findings align with previous work showing that seagrass meadows offer superior refuge to the smallest and most vulnerable size classes of juveniles (e.g., Hovel and Lipcius 2001; 2002, Orth and van Montfrans 2002, Johnston and Lipcius 2012, Bromilow and Lipcius 2017, Shakeri et al. 2020). The substantially higher survival in SME relative to sand further supports experimental results indicating that salt marsh structure can provide at least partial refuge compared to unstructured habitats (Johnston and Caretti 2017, Miller et al. 2023).

Importantly, the refuge value of SME increased sharply with size. By 15 mm CW, survival in SME and seagrass was comparable, and by 33 mm CW, SME survival was equal to that of seagrass. In contrast, survival in seagrass remained high but relatively size-invariant, consistent with previous work (Johnston and Lipcius 2012). This interaction between size and habitat suggests that the protective value of habitat structure depends not only on structural complexity but also on the size-specific vulnerability of juvenile crabs to predation.

This size-dependent pattern likely arises from how structural complexity mediates refuge quality across different predator and prey sizes. For example, Orth and van Montfrans (2002) found that salt marsh shoots did not protect small juveniles (2.1 – 12.6 mm CW) from predation by mummichogs (*Fundulus heteroclitus*), but juveniles reached a size refuge from these smaller predators by 9 – 12 mm CW. Our results suggest that while early-stage juveniles in SME remain vulnerable to small predators capable of maneuvering through interstitial spaces between marsh shoots, larger juveniles evade predation by exceeding the gape sizes of these predators (Urban 2007). At the same time, the structure of the marsh still impedes foraging by larger predators – such as adult conspecifics (e.g., Johnston and Caretti 2017, Miller et al. 2023) – thus enhancing refuge with size. Together, these patterns imply that both seagrass and SME habitats provide meaningful, albeit size-dependent, refuge from predation, with unstructured sand offering the least protection across sizes.

Using average growth rates from earlier work, the trade-off between growth and mortality risk appears to initially favor seagrass meadows for early-stage juveniles, but increasingly shifts toward upriver salt marsh habitat as crabs grow. This pattern mirrors recent findings that documented fine-scale (5 mm CW) ontogenetic shifts in habitat use, where the smallest juveniles ( *≤* 15 mm) were most abundant in seagrass meadows, while juveniles between 16 – 25 mm were progressively more common in salt marshes, particularly in areas with high turbidity (Hyman et al. 2022; 2024b). This shift possibly reflects both increased access to prey and reduced predation risk as individuals outgrow smaller predators. Larger juveniles are strongly associated with bivalve prey commonly found in estuarine turbidity maxima (Seitz et al. 2003; 2005), and transitions from seagrass to unstructured or marsh-dominated upriver habitats have long been hypothesized as a consequence of the growth–mortality trade-off (Lipcius et al. 2005). Importantly, salt marshes occur not only upriver but also in turbid mid-estuarine locations throughout the York River and Chesapeake Bay, and larger juveniles are consistently associated with areas that combine high turbidity and extensive marsh habitat (Hyman et al. 2022). Taken together, our findings suggest that turbid, structurally complex salt marshes function as intermediate nurseries for juveniles that have attained a partial size refuge from smaller predators, but still benefit from the combined refuge and enhanced food resources that these habitats offer.

Placed within the context of this previous work, our results indicate that the current paradigm of juvenile blue crab ontogenetic habitat utilization requires revision. We propose a revised conceptual model describing differences in juvenile blue crab habitat use at different size classes (Fig. 4). Ingressing megalopae predominantly settle into seagrass beds or other SAV until *∼* 15 mm CW (Orth and van Montfrans 1987, Hyman et al. 2024b, Johnston and Lipcius 2012), although small juveniles may utilize salt marsh vegetation as an alternative nursery if they do not encounter downriver SAV as megalopae (Stockhausen and Lipcius 2003) or after emigrating from seagrass to avoid adverse density-dependent effects (Etherington and Eggleston 2000, Etherington et al. 2003, Reyns and Eggleston 2004, Blackmon and Eggleston 2001). Once juveniles reach *∼* 15 mm CW, they increasingly disperse to SME and tidal marsh creek habitats, where greater food availability supports higher growth rates (Seitz et al. 2005), and salt marsh vegetation provides refugia from predation, especially during vulnerable molting periods (Ryer et al. 1997, Lipcius et al. 2007, Hines 2007). The higher food availability in upriver salt marsh habitat near the estuarine turbidity maximum appears particularly valuable for juveniles *≥* 20 mm (Hyman et al. 2022). By 35 mm CW, juveniles increasingly reside in turbid salt marsh or adjacent unstructured habitat (Lipcius et al. 2005, Hyman et al. 2022).

**Figure 4.**
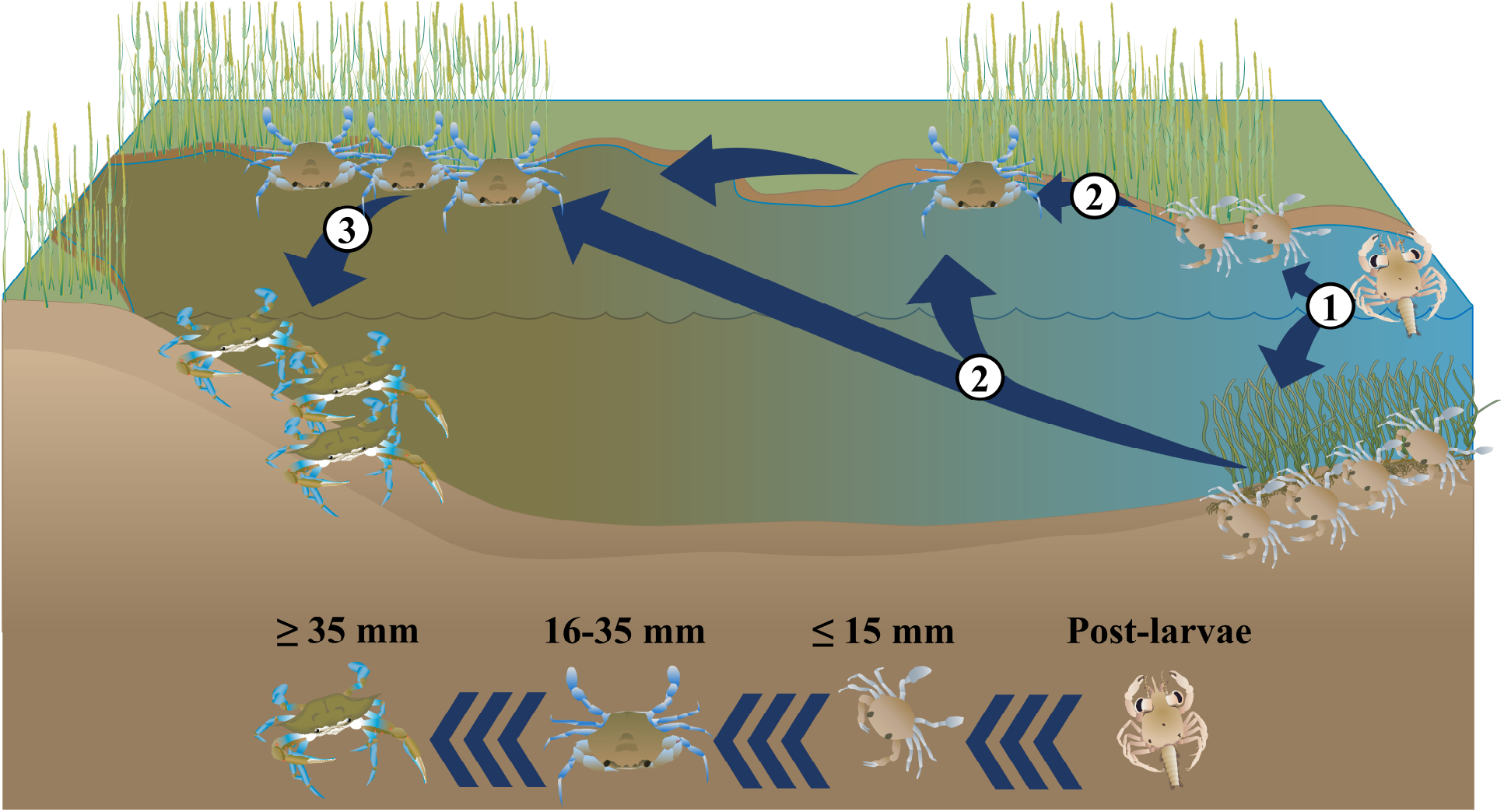
Conceptual diagram illustrating revised ontogenetic habitat shifts of juvenile blue crabs in mid-Atlantic estuaries. Numbered arrows indicate transitions between habitats as crabs grow: (1) Ingressing post-larvae initially settle in seagrass beds or, if unavailable, adjacent salt marsh habitat; (2) after *∼* 15 mm carapace width (CW), juveniles increasingly emigrate to turbid, structurally complex salt marsh and tidal creek habitats to exploit higher food availability and intermediate structural refuge;(3) after *∼* 35 mm CW, juveniles increasingly occupy turbid, unstructured habitats as size provides sufficient refuge from predation. Graphics were obtained in part through the [Integrated Application Network (IAN) Image Library](https://ian.umces.edu/media-library/).

### Relevance

Our results expand upon previous work documenting patterns of habitat utilization in juvenile blue crabs (Seitz et al. 2005, Lipcius et al. 2005, Hyman et al. 2022; 2024b) to provide a plausible mechanism – trade-offs between growth and survival – underlying ontogenetic habitat shifts in mid-Atlantic estuaries. Our findings agree with shifts in habitat utilization from seagrass beds but emphasize the role of structured salt marsh habitat as a critical nursery for juveniles within intermediate size ranges before reaching a complete size refuge from predation.

Our results also add to a growing body of research highlighting a need to preserve the complete chain of nursery habitats used by juvenile blue crabs (i.e., seagrass meadows and salt marshes) before entering adult habitats (e.g. Johnson and Eggleston 2010, Hyman et al. 2022; 2024b). Seagrass and salt marsh declines have received considerable attention in Chesapeake Bay (e.g. Moore et al. 2014, Schieder et al. 2018). Eelgrass (*Zostera marina*) beds are declining due to direct and indirect anthropogenic influences such as land-use change and long-term warming of Chesapeake Bay (Orth et al. 2010, Moore et al. 2014, Patrick et al. 2018, Hensel et al. 2023). Similarly, salt marshes have been reduced by coastal development and shoreline hardening (Silliman et al. 2009) as well as sea level rise (Kirwan and Megonigal 2013, Schieder et al. 2018). Although losses in blue crab secondary production associated with seagrass declines have received considerable attention (e.g. Hovel and Lipcius 2001; 2002, Mizerek et al. 2011, Ralph et al. 2013, Johnston and Lipcius 2012), the effects of salt marsh loss on blue crab population dynamics remain a major data gap for Chesapeake Bay and other mid-Atlantic estuaries. Moreover, ratios of marsh-migration to marsh-erosion associated with sea level rise are spatiotemporally variable, and higher rates of salt marsh loss are expected in upriver regions such as within the York River (Schieder et al. 2018), which are potentially the most valuable for later-stage juvenile blue crabs. Additional empirical and mechanistic modeling is required to quantify how shifting seagrass and salt marsh distributions, as well as novel nursery habitats such as algal patches, will affect blue crab population dynamics both within Chesapeake Bay and among Northwestern Atlantic estuaries.

### Caveats and limitations

While this study offers valuable insights into the ontogenetic mechanisms underlying blue crab habitat shifts, a key caveat is the potential confounding influence of historically low adult blue crab abundance on habitat-specific juvenile survival—particularly in SME. We observed higher juvenile survival in SME than previously reported in other systems (Shakeri et al. 2020). This discrepancy may be partly explained by the unusually low abundance of adult male blue crabs during our study period, as 2021 saw the lowest adult male population levels recorded since the onset of fisheries-independent monitoring in 1990 (MDNR 2019, CBSAC 2022). Additionally, this discrepancy may have been in part due to differences in methodology (e.g., tether length) or predator suite. Given that adult blue crabs are significant predators of juveniles (Moody 2003, Eggleston et al. 2005, Hines 2007, Lipcius et al. 2007, Bromilow and Lipcius 2017), this reduction in adult density likely reduced predation pressure. Therefore, elevated survival rates in SME habitats may partly reflect temporarily reduced adult predation rather than inherent habitat quality, potentially inflating our observations of SME’s relative refuge value. We advise caution in generalizing these results and recommend future replication during periods of higher adult abundance to evaluate the robustness of these patterns. We also urge further investigation to determine whether this pattern persists or if our results were influenced by local and/or ephemeral processes. Additional research in this area is crucial to validate and understand these results more comprehensively.

## Acknowledgment

ACH thanks D. Eggleston and C. Patrick for their ideas as members of ACH’s PhD Committee, J. Shields for comments on earlier versions of this manuscript, and C. Miller for assistance with tethering work in the field. Funding for the research and manuscript preparation was provided by the Willard A. Van Engel Fellowship to ACH, National Science Foundation grants EEID 2207343 to J. Shields, R. Lipcius, and K. Reece, and OCE (REU) 2243957 to R. Seitz and G. Massey, and the joint National Marine Fisheries Service/Sea Grant Population and Ecosystem Dynamics Graduate Research Fellowship to ACH.

